# Strong and Fluctuating Sequence Constraints Drive *Alu_* Evolution

**DOI:** 10.1101/094326

**Authors:** Aaron C. Wacholder, David D. Pollock

## Abstract

Though *Alu* elements are the most common and well-studied transposable elements in the primate genome, *Alu* evolutionary dynamics remain poorly understood. To better understand these dynamics, we improved our recently introduced Bayesian transposable element ancestral reconstruction method to incorporate automated alignment and be more computationally efficient. We then used it to reconstruct the relationships among almost 800,000 *Alu* elements in the human genome. We identified the phylogenetic network relating 154 ancestral replicative *Alu* sequences, and found that the aligned ancestors vary at only 56 out of ~300 sites. We show that the limited number of variable sites among replicative *Alu* ancestors is best explained by strong sequence constraints on *Alu* replicative capacity. Moreover, the pattern of variation suggests that sequence constraints fluctuated over the course of *Alu* evolution, driving the extinction of older *Alu* subfamilies and the birth of newer ones. Previous analyses have taken the tight clustering of *Alu* sequences with age as evidence that all *Alu* sequences are descended from a small number of "master elements." Our results imply instead that the clustering of *Alu* sequences with age results from fluctuating sequence constraints, and that there were over 4,000 replicative loci during the course of *Alu* evolution, most of which were disabled by mutation before mutating to new replicative sequences. We also predict which sites have been functionally important for replication, and how these sites have changed over time. The newly clarified dynamics of *Alu* evolution invalidate assumptions used in common methods of transposable element classification and phylogenetics.

**Significance Statement:** Transposable elements are genomic sequences that can insert copies of themselves elsewhere in the genome. *Alu* is the most abundant transposable element in primates, making up 10% of the human genome. Due to its ubiquity and tendency to cause genomic instability, *Alu* has played a major role in shaping primate genomes. Characterizing the trajectory of *Alu* evolution is important for understanding how the human genome evolved.

Previous analyses of *Alu* concluded that a tiny number of elements generated all copies, and existing classifications of *Alu* reflect that conclusion. In a whole-genome analysis, we determine that many more elements were replicative than previously understood, indicating that current classifications of *Alu* are flawed. We develop an alternative reconstruction of *Alu* evolutionary history.

## Introduction

Transposable elements (TEs) are genomic sequences that can insert copies of themselves elsewhere in the genome. Transposable elements and TE remnants comprise a large portion of many eukaryotic genomes (1), including a majority of the human genome (2). Understanding TE evolution is therefore vital to understanding how genomes evolve. Yet many fundamental aspects of TE evolutionary dynamics remain poorly understood.

Of key importance is to determine the conditions under which a TE is capable of replication. The constraints on TE replication can be divided into two general categories: locus and sequence. Sequence constraints include any factor relating to the primary sequence of the element, while locus constraints include factors relating to the position of the element in the genome, such as local chromatin structure or the presence of nearby regulatory elements. The relative importance of locus and sequence constraints determine many features of TE dynamics. If locus constraints are strong and sequence constraints weak, most TE insertions will land in inhospitable regions and never replicate. The TEs that do land in favorable regions may last a long time, producing many copies in their lifespan, as most mutations will not eliminate replicability. In contrast, if sequence constraints are strong and locus constraints weak, a large portion of elements will land in favorable regions and therefore be capable of replication at time of insertion. Individual elements will be short-lived, however, as mutation will tend to degrade replicative capacity.

Early research on human LINE and SINE TEs suggested that these elements evolved under extreme locus constraints and weak sequence constraints (3–5). According to the early Master Element Model of TE evolution, a single *Alu* "master element" located at a favorable locus produced all of the over 1 million *Alu* copies in the human genome. Evidence for this model came from the observation that clusters of Alu elements of similar age all appeared to be descended from a single source sequence; it was inferred that each cluster corresponded to a particular sequence at the master locus, with transitions between clusters caused by mutation at that locus. Subsequent research has revealed that this model is not precisely correct. Using whole-genome *Alu* data, Price et al. estimated that at least 143 replicative elements existed during the course of *Alu* evolution (6), and human polymorphism data indicates the existence of at least 31 currently-active *Alu* elements (7). In response to the revelation of multiple source elements, modifications were proposed to the master model to allow for the possibility of a small number of concurrently-active source elements (8,9). However, these new models also assume that only a tiny fraction of *Alu* loci are compatible with replication, and that the few elements at these favorable locations are the origin of most *Alu* elements in primate genomes.

Most previous analyses of *Alu* evolutionary dynamics were based on small subsets of *Alu* elements. To better characterize *Alu* dynamics, we improved our Bayesian TE ancestral reconstruction method (AnTE) to simultaneously determine the evolutionary relationships among almost 800,000 *Alu* loci in the human genome. We previously demonstrated, using a selected subset of *Alu* sequences, that our AnTE method can provide a higher-resolution picture of TE relationships than the popular CoSeg method (10). Our new method, AnTE2, improves on AnTE by enabling the analysis of evolutionary relationships among a much larger number of TE sequences. Applying AnTE2 to *Alu*, we identified many previously-unidentified *Alu* ancestral sequences, which enabled more precise estimation of sequence and locus constraints on *Alu* replication.

Surprisingly, and in sharp contrast to previous models, we find that *Alu* evolution is characterized by strong sequence constraints but relatively weak locus constraints. We propose that in each period of *Alu* evolution, replicative capacity was constrained to elements within a small region of sequence-space. This hypothesis, which is consistent with experimental evidence (11,12), explains why *Alu* elements with similar ages appear to be descended from similar ancestral sequences, but allows for the possibility that these ancestral sequences were shared by a large number of historically-replicative loci. Indeed, we find evidence that although there were only hundreds of replicative sequences, there were potentially thousands of replicative loci.

## Results

### A Network of *Alu* elements

The AnTE2 algorithm identifies a set of ancestral sequences from a dataset of TE elements. We define "ancestral sequence" as an ordered set of nucleotides that replicated at least once in the course of *Alu* evolution. As there is some ambiguity to the term “sequence” as used in molecular biology, it is important to note that, as we use it, the term “ancestral sequence” refers to the sequence itself and not any particular locus. Thus, a single ancestral sequence can correspond to any number of identical replicative loci. The AnTE2 algorithm also estimates, for each identified ancestral sequence, the total number of copies that were replicated from loci with that sequence and the average age of the copies of that sequence.

We applied AnTE2 to 779,310 *Alu* elements from the hg38 assembly of the human genome. AnTE2 identified 154 high-confidence ancestral *Alu* sequences, which differ at 57 sites (Supplementary Table 1). A total of 63 non-consensus variants were identified among ancestral sequences, including 62 single nucleotide variants and a single 2-base pair indel variant (Supplementary Table 2). We infer that most ancestral sequences produced relatively few copies, while a minority were responsible for most of the elements in the genome (Supplementary Figure 1). For example, the ten most productive ancestral sequences out of 154 were responsible for a combined 62% of elements, and the top thirty produced 83% of elements.

We visualized relationships among the replicative ancestors using a network structure in which each node represents an ancestral sequence and edges were drawn between nodes that differ by a single variant; we refer to this structure as a *replicative network* (Supplementary Figure 2). The network is highly connected overall, with mean degree 2.76. Visual inspection of the *Alu* replicative network indicated that there are four major clusters in addition to 15 nodes outside any major cluster. These clusters are similar to the traditional grouping of *Alu* sequences into subfamilies *AluJ, AluS*, and *AluY*, but the *AluS* subfamily is split into two clusters, which we term *AluS1* and *AluS2* (Table 1). To clarify the relationship among the major clusters, we added a second set of thinner edges between all pairs of unconnected nodes that differ at two positions (Figure 1). By coloring the replicative ancestral sequence nodes based on their estimated ages, it can be seen that the major clusters are arranged sequentially in time, with each cluster representing a distinct time period of *Alu* evolution.

**Table 1.**

| Cluster | Mean age | Node count | Variant count | Element count | Replication-compatible variants (95% CI) |
| --- | --- | --- | --- | --- | --- |
| AluY | 0.56 | 25 | 18 | 114,025 | 20-40 |
| AluS2 | 0.89 | 21 | 9 | 53,220 | 9-14 |
| AluS1 | 0.9 | 46 | 18 | 468,185 | 18-23 |
| AluJ | 1.47 | 62 | 28 | 143,795 | 29-39 |
| Entire Network | 1.08 | 154 | 63 | 779,310 | 66-79 |

**Figure 1.**
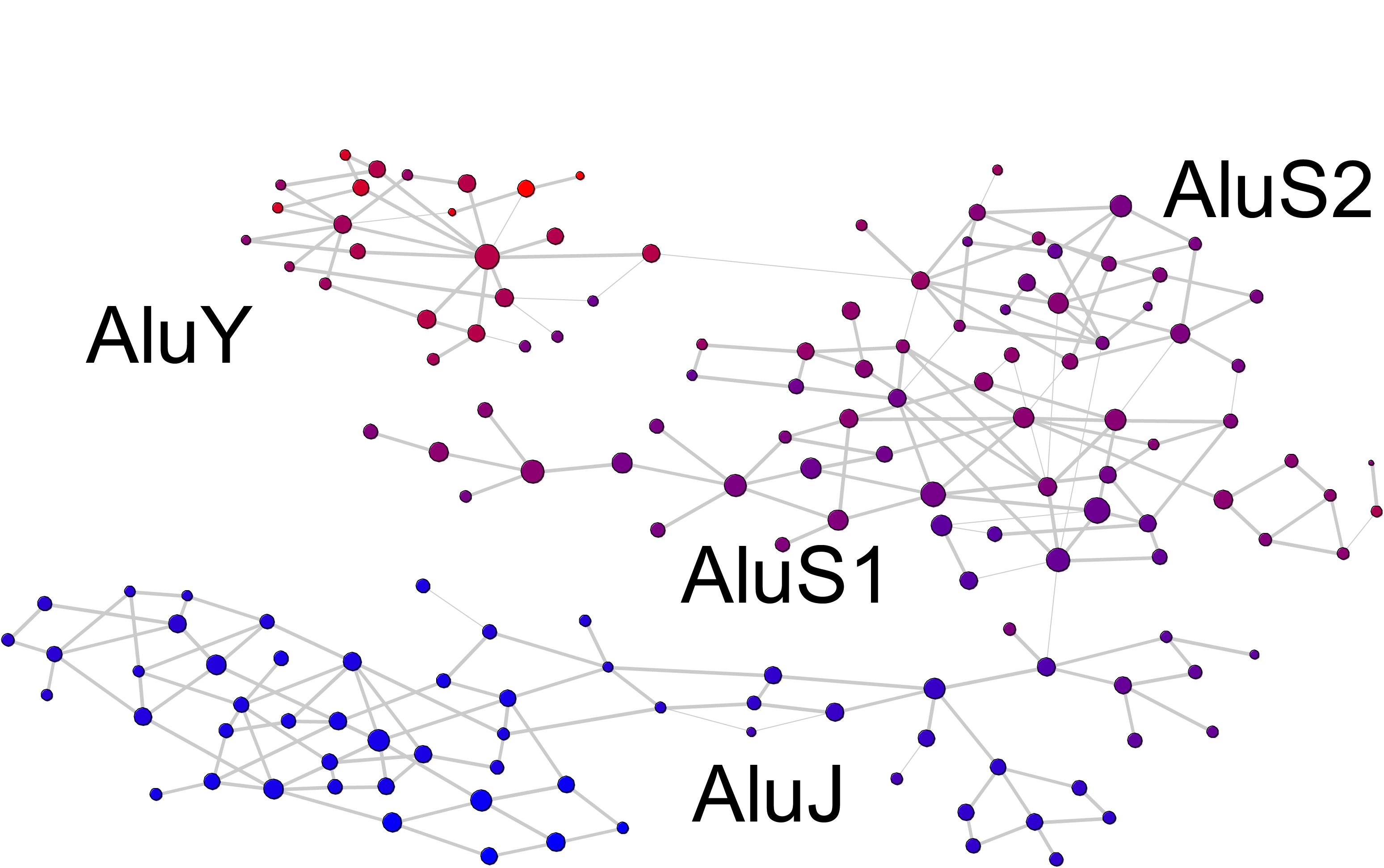
*Alu* Replicative Network. Nodes in the network represent ancestral sequences. Thick edges connect nodes differing by a single variant; thin edges connect nodes differing by two variants, if no path consisting of single variants exists between them. Node size is proportional to the log of the replicative output of the ancestral sequence. Node color corresponds to age, with a redder shade indicating a younger ancestral sequence.

### Most Sites in *Alu* are Constrained

The AnTE2 method identified 63 non-consensus variants among ancestral sequences out of 396 variants that could possibly distinguish *Alu* ancestral sequences in our analysis. We do not expect this to be an exhaustive set, as low-output replicative sequences are difficult or impossible to detect, and we conservatively eliminated sequences with low evidence for having been ancestral. However, as all other variants exist at frequencies similar to expectation from substitution rates (Supplementary Figure 3), any missed variant that was present in replicative sequences must have existed only in replicative sequences with very low output.

The observation that the number of variants in ancestral sequences, 63, is much smaller than the number of ancestral sequences, 154, indicates that the variants that do appear within ancestral sequences often distinguish multiple pairs of ancestral sequences. However, most possible sequence variants (86%) never appear in even one ancestral sequence. That some variants appear to be involved many times in the formation of new ancestral sequences and others not at all suggests that *Alu* replication is constrained to some degree by sequence. We use simulation based on the replicative network to evAluate this hypothesis.

Consider the generation of a novel ancestral sequence by mutation at a replicative locus with an older ancestral sequence. If TE replication was completely unconstrained by sequence, and the mutation rate was equal at all sites, then all sites would be equally likely to be involved in the transition from one ancestral sequence to the next. We could then simulate the sites involved in each transition by simply randomly sampling from all possible sites. We do this sampling 1000 times for each of the 153 transitions to new ancestral sequences that would have been necessary to produce all 154 ancestral sequences identified by AnTE2. Among all 1000 simulations, between 88 and 103 total sites are involved in the generation of new ancestral sequences, far more than the actual 57 varying sites identified by AnTE2. This indicates that not all sites are equally likely to be involved in the generation of a new ancestral sequence, which could be a consequence of either differing mutation rates or sequence constraint. Next, we repeat the analysis accounting for the approximately two-fold difference in mutation rate between transitions and transversions. In 1000 resamplings, between 85 and 108 total sites are involved in transitions to new ancestral sequences, still far higher than the 57 identified by AnTE2. We conclude that *Alu* replication is constrained by sequence.

We next attempt to quantify the degree of sequence constraint in *Alu* using a similar strategy. We resample the sites involved in the generation of ancestral sequences assuming that *N* sites chosen at random completely eliminate replicative capacity (and thus cannot be sampled) and the remainder have no effect on replicability at any point in time. We resample 1000 times for each *N*, determining which *N* have the observed number of sites differing among ancestral sequences, 57, within their 95% confidence interval. This analysis indicates that the number of constrained sites is between 86 and 97, 55-62% of the total 156 sites considered.

One source of sequence constraint in *Alu* comes from the need for folding into an appropriate structure for retrotransposition. The consensus Alu sequence contains 162 base-paired positions and 120 single-stranded positions (11) and replication-compatible variants identified by AnTE2 appear 66% more frequently in single-stranded positions (p=.033, Fisher exact test). Of the 28 variants in paired positions, six vary between the canonical G-C and wobble G-U base-pairing, and eight are involved in four pairs in which both sides of the base-pair vary, allowing at least one canonical base-pairing for each variant. Interestingly, the transition between *AluS1* and *AluS2* and the transition between *AluS2* and *AluY* each involve a pair of variants that are base-paired to each other, while the transition from *Alu*J to *AluS1* involves a pair of variants that together are associated with an alternative conformation at a site of Alu-SRP binding (11). In each of these three cases, we identify no *Alu* ancestral sequences which contain only one of the non-consensus variants. Thus, it appears that each of the transitions between major *Alu* classes is associated with a pair of interacting variants in which both variants are required for detectable rates of replicability.

### Changing Sequence Constraint over the *Alu* Network

In the above analysis, we estimated the number of constrained sites in *Alu* under the assumption that sequences constraints were constant in *Alu* evolutionary history. To determine whether this assumption is justified, we investigated whether sequence constraints change over time using the structure of the replicative network.

The abundance of cycles of size four in the replicative network reflects patterns of sequence constraint. To understand this relationship, consider a hypothetical replicative network consisting of three ancestral sequences *A*, *B*, and *C*, in which *B* differs from *A* by a single variant at site *x* and *C* differs from *B* by a single variant at a distinct site *y* (Figure 2). This is represented by a linear replicative network in which *A* shares an edge with *B* and *B* shares an edge with both *A* and *C*. Suppose that ancestral sequence *C* experiences a mutation that generates a new replicative sequence *D*. If this mutation is a reversion to the variant A has at position x, then the resulting replicative network will form a cycle. Any other mutation would result in a linear structure. The probability a cycle is formed, then, depends on the relative probability of the one possible cycle-forming mutation compared to the probability of all other mutations that could produce a novel ancestral sequence. If sequence constraints are weak, then there would be many mutations that would produce new ancestral sequences, so it would be improbable that the generation of any particular novel ancestral sequence would result in a cycle in the replicative network. Alternatively, if sequence constraints are strong, then the cycle-forming ancestral sequence would be one of a small number of possibilities, and would thus be relatively likely. Cycles in the replicative network result from convergence and reversion events, and the frequency of such events depend on the strength of sequence constraint.

**Figure 2.**
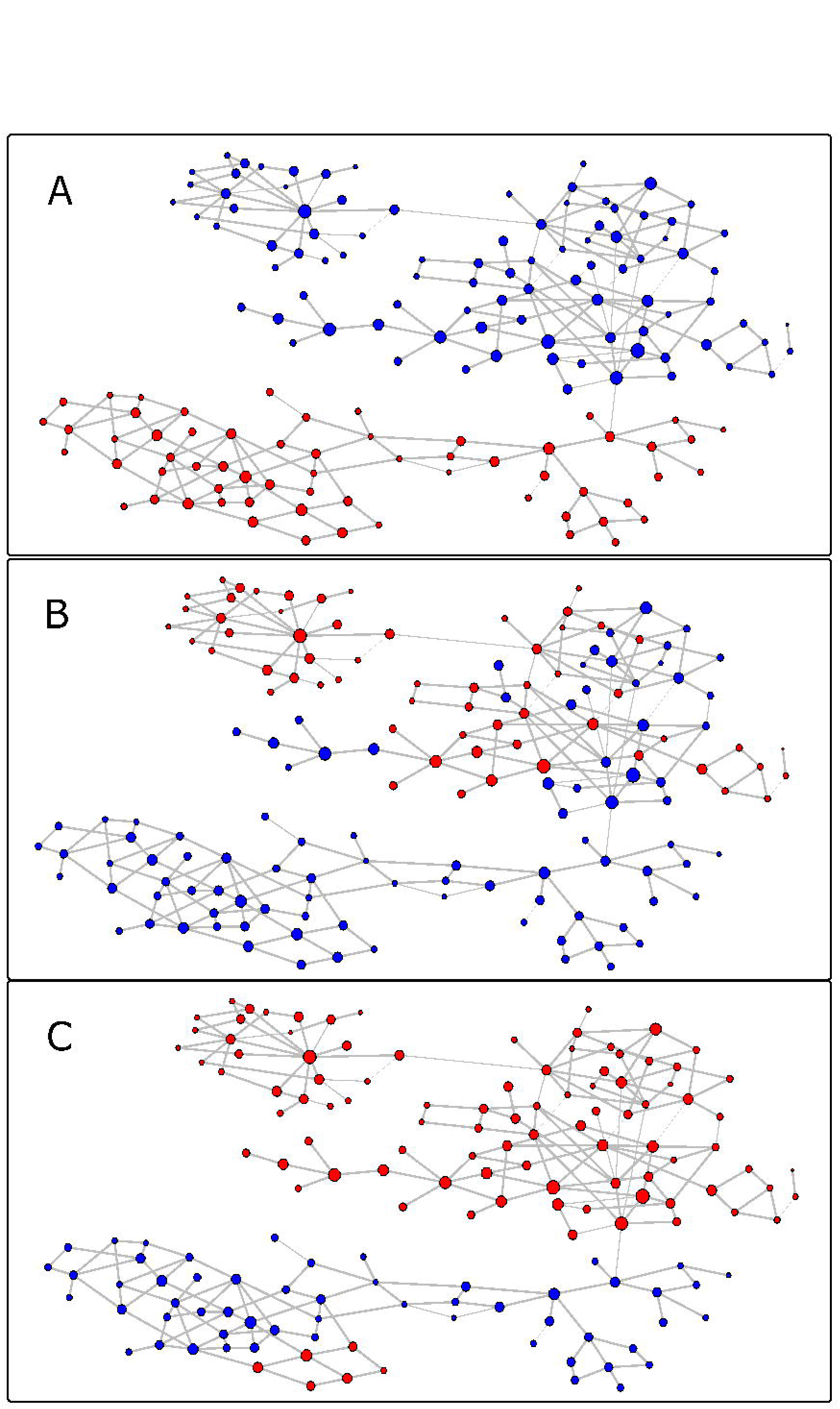
Variant Patterns. Replicative network as in Figure 1, with nodes colored based on which variant exists at a position. Combinations with the rarer variant are colored in red; those with the more common variant are colored blue. A) Position 94; C in red, G in blue B) Position 153; G in red, C in blue C) Position 70; A in red, G in blue.

We used the abundance of cycles on the replicative network to estimate the number of variants compatible with replication for the average ancestral sequence on the replicative network. Though the details are somewhat complex (see Appendix), we essentially compare how often cycle-forming events must have occurred to obtain the network structure observed to the frequency such events would be expected to occur under different scenarios of sequence constraint. Of 153 events generating new ancestral sequences, 74 (.48) resulted in the formation of cycles in the replicative network. If all variants identified as existing in at least one replicative sequence were compatible with replication over the entire history of *Alu*, then no more than 21% of events generating new ancestral sequences would be expected to form cycles. The probability of 74 or more events out of 153 forming cycles with a 21% chance of each event forming a cycle is 1.48e-14; thus we conclude that some variants that were compatible with replication at one point in time were not compatible over the entire course of Alu evolution. We estimate that, for the average ancestral sequence, no more than 9.8% of possible variants (38.8 of 396 considered) would be compatible with replication. As this is substantially smaller than the 63 variants identified by AnTE2 as existing in at least one time period of *Alu* evolution, we infer that sequence constraints changed substantially during *Alu* evolution.

Patterns of variation at individual sites offer further clues as to how sequence constraints have changed through *Alu* evolution (Figure 3, Supplementary Figure 4). Some sites, such as position 94, exhibit sharp transitions on the network (Figure 3A). All replicative ancestors in the oldest component appear to have a C at this position, while all younger ancestors have a G. Other sites, such as position 153 (Figure 3B) or position 70 (Figure 3C), have more complicated patterns, with one variant dominant in some regions of the network and both variants coexisting in others.

**Figure 3.**
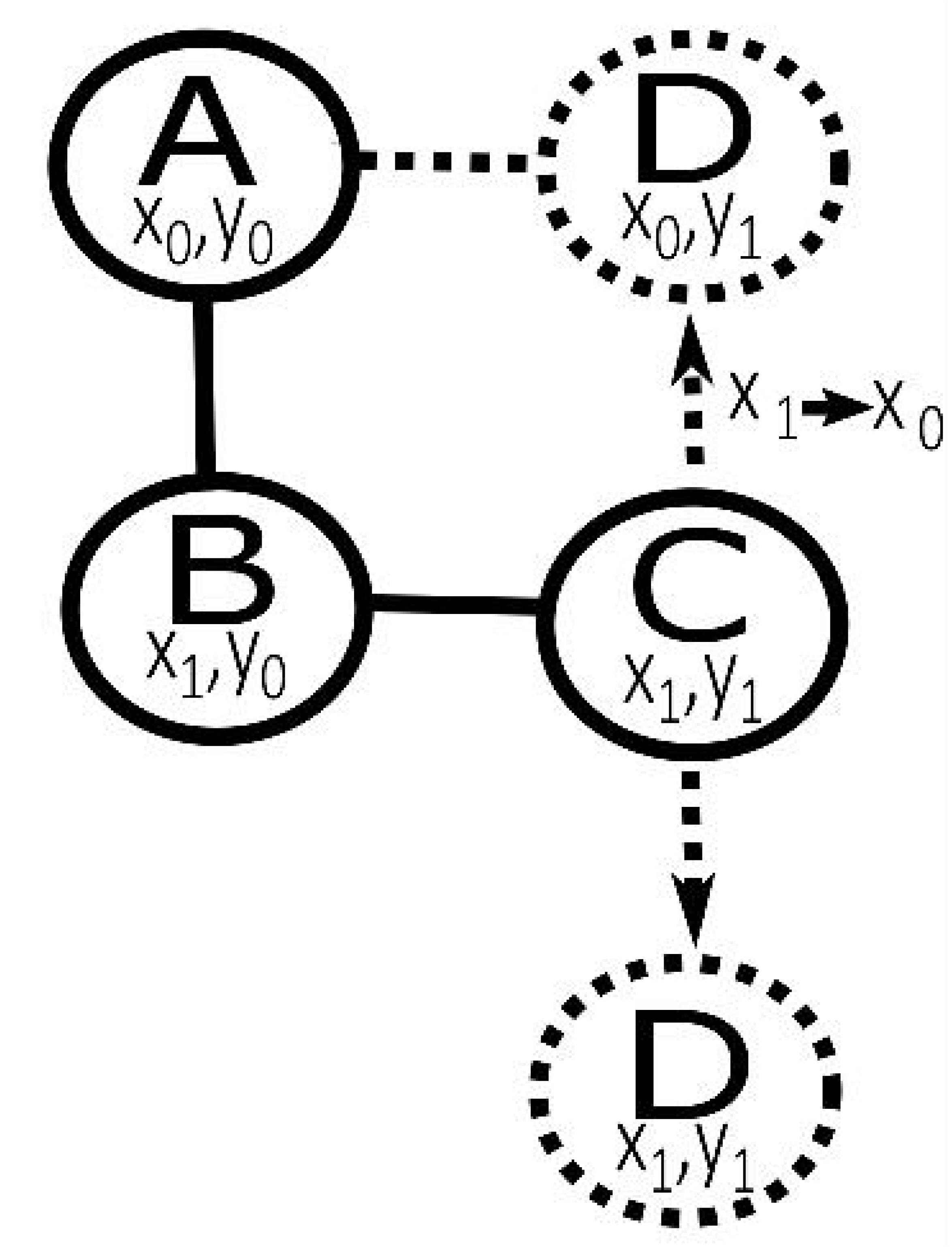
Producing Cycles in the Replicative Network. The nodes *A, B*, and *C* represent nodes in a hypothetical replicative network that vary at positions *x* and *y*. The variants at position *x* and *y* are listed below the node label. Edges connect nodes differing by a single variant. If a new replicative ancestor is produced by ancestor *C*, a cycle will be created in the replicative network only if the mutation distinguishing that ancestor is from *x*_1_ to *x*_0_; any other new ancestral sequence generated from *C* will maintain the replicative structure.

### Estimating the Number of Replicative Loci

The AnTE2 algorithm provides estimates for the total number of ancestral sequences, but it is not straightforward to infer from this the number of replicative loci. Though most previous research on *Alu* has assumed that each ancestral sequence corresponds primarily to a single replicative locus (6), this need not be the case: a single ancestral sequence might have been represented by many identical replicative loci during its period of activity.

The stronger sequence constraints are, the weaker locus constraints must be for a TE family to maintain viability. If, as suggested by the analysis above, only around 9.8% of mutations are compatible with continuing replication, then for each mutation occurring in a replicative locus leading to a new ancestral sequence, we can infer the existence of around nine mutations which instead disable previously-replicative loci. As the 154 identified ancestral sequences require 153 events creating the first replicative loci with that sequence, this implies the existence of around 1386 (9 * 153) loci disabled by mutation, which means at least that many loci were replicative at some point.

Further evidence for many replicative loci comes from the frequency of convergence across the replicative network: because few mutations would be convergent, a high frequency of convergence suggests the existence of many replicative loci in which convergent mutations could potentially occur. The clearest demonstration of this principle is the 2 bp deletion variant at position 65-66. The rate at which this deletion mutation occurs can be estimated by observing its frequency in *Alu*J sequences, whose ancestral sequences do not contain this deletion mutation. Only 110 of 143,795 *AluJ* elements contain this variant, giving a rate of 0.08%, less than 1/10^th^ the rate of any single nucleotide mutation. For *AluS* and *AluY* sequences, in which this deletion variant is abundant, it is therefore likely that most sequences containing the variant descended from ancestral sequences that already contained the variant and did not obtain it through independent mutations. The evolutionary trajectory of ancestral sequences with the deletion variant should thus have been largely independent of the trajectory of ancestral sequences without the variant. Despite this, there is large agreement between these trajectories, indicating a great deal of completely convergent evolution. In the three youngest clusters there are 54 ancestors with the deletion variant and 38 without; 27 of these are pairs that match at all other positions except the deletion variant, meaning that exclusive of the deletion variant, a majority of the sequences are convergent (50% of those with the deletion and 71% of those without). Though it is not simple to precisely estimate a count of replicative loci from these data, it does suggest that the evolutionary trajectory of *Alu* evolution was to a large extent controlled by sequence constraints rather than loci constraints.

## Discussion

We identified 154 high-confidence *Alu* ancestral sequences using a novel method for TE ancestral reconstruction that avoids restrictive assumptions made by other methods. This method, AnTE2, improves on our previous AnTE algorithm by employing an iterative approach to greatly increase the scale at which ancestral reconstruction is possible and thus allow us to analyze nearly one million sequences simultaneously. By identifying numerous new ancestral sequences, we were able to determine that *Alu* replication was characterized by strong sequence constraints that changed over time. Contrary to the typical assumption that each ancestral sequence corresponds to a single loci (6), we find strong evidence that there were over a thousand replicating ancestral loci and that ancestral sequences were mostly represented by multiple replicative loci.

### The Network Model of *Alu* Evolution

As with these previous studies (3,13,14), we find that the number of identified replicative sequences, 154, is very small relative to the number of elements in our dataset (779,310). Previous studies interpreted this discrepancy as evidence that few loci are capable of replication. Our findings support an alternative explanation: there are few ancestral sequences because most sequences are incompatible with replication, with most ancestral sequences likely represented by a large number of loci.

The key evidence that *Alu* evolution was affected by sequence more than locus constraints comes from the observation that, of the variants present in at least one ancestral sequence, many are present in a large number of ancestral sequences, while most variants never appear among replicative sequences. This feature of *Alu* evolution is only readily apparent with the identification of a sufficiently large sample of *Alu* ancestral sequences. It is understandable, then, that it went unnoticed by many previous analyses, which were based on relatively small subsets of extant elements, and therefore missed many low-output ancestral sequences. The only previous study of *Alu* evolutionary relationships that analyzed a large fraction of the elements in the human genome was conducted by Price et al (6). However, the CoSeg method used by these authors (15), which is based on iterative splitting of subfamilies, forbids splitting off a new subfamily on the basis of a variant that was already used to split a subfamily earlier in the process (10). Thus, because of its design their analysis was unable to identify the highly-networked structure we find in *Alu*, in which a single variant can differentiate many pairs of ancestors.

We propose a new model of TE evolution to account for our results, which we label "the Network Model." In this model, TE replication is strongly dependent on sequence, with most mutations eliminating or severely reducing replicability. An element with a sequence close to optimal at a favorable locus will tend to produce new replicative copies at other locations in the genome, growing the pool of identical elements. In time, replicative elements mutate. Most mutations permanently inactivate the element, while a minority result in transfers from one pool of replicative elements to another.

We call this the "network model" because if edges are drawn between pools of ancestral sequences that differ by a single variant, a highly-connected network structure emerges. This structure results from convergent mutation and reversion between the small number of variants that permit replication. If a pair of sequences differing at one site mutate at another site to the same replication-compatible variant, the four sequences involved form a cycle in the replicative network representing each possible combination of the two variants. The network model thus explains the highly connected replicative network generated by AnTE2 as well as the long-noted observation that *Alu* replicative sequences contain only limited sequence variation.

### The Replicative Network

Early studies of human TEs classified elements into groups called "subfamilies" (3, 4,16, 17). Under the original master element model, a single locus produced all elements in the genome, maintaining replicability through numerous substitutions. In this model, a subfamily referred to the set of elements descended from the master locus when it had a particular sequence. The trajectory of a TE family evolving under a master model could then be represented by a linear progression between subfamilies, with transitions between subfamilies occurring as a result of substitution events at the master locus.

As new research identified multiple replicative elements, it became clear that *Alu* evolution could not be represented by a linear progression of subfamilies (13,18). Under the new understanding of *Alu* evolution, it was possible for a replicative element to produce a new replicative element; this allowed the evolution trajectory of *Alu* to branch. Thus, later authors represented *Alu* evolution using a directed tree of subfamilies. Though the term "subfamily" has rarely been precisely defined in the human TE literature, in practice subfamilies were identified by clustering extant elements by sequence similarity (15).

In light of our results, there are two major problems with the "tree of subfamilies" representation of *Alu* evolution. First, due to the possibility of homoplasy, sequence similarity need not indicate close relatedness. We find numerous ancestral sequences that are very similar to other ancestral sequences and evidence for frequent convergent evolution. It is thus not improbable that convergent evolution produced two identical replicative elements twice by distinct pathways from different source sequences. The descendants of these two identical replicators would be the same at insertion, and thus group together if clustered by similarity, but would be more closely related to elements in other clusters than to each other. Moreover, as the "subfamily" identified by clustering would have two ancestors, which is forbidden in a tree, no tree could accurately describe its evolutionary relationship to its ancestral subfamilies. Secondly, even if the evolution of *Alu* could be accurately described by a tree, the close similarity of many ancestral sequences means that many possible tees are similarly likely. Picking any one tree would poorly represent the space of plausible paths.

For these reasons, we believe that the evolution of *Alu*, and perhaps other TE families with similar dynamics, are best represented by replicative networks. Instead of subfamilies, nodes in the replicative network represent ancestral sequences; i.e., a particular ordered set of bases. Each ancestral sequence is associated with an estimated count of elements that were replicated from that sequence, but no claim is made about the evolutionary relationship among those elements. For ease of visualization, undirected edges are drawn between nodes which differ by a single variant. Thus, the most likely paths by which a replicative element could transition by mutation to a different ancestral sequence is readily seen.

### Explaining Patterns of Alu Sequence Constraint

The network model implies that the pattern of ancestral *Alu* sequences can provide potentially useful predictions about the effect of variation on replicative capacity. In a master element model, most replication-compatible variants are never reached by mutation at replication-compatible loci, so the observed evolutionary trajectory of a TE family gives little information about the effect of most variants on replicability. In the network model, because there are many replicative loci there are many opportunities for a TE family to sample the effect of mutations on replicability. Therefore, the observation that a site never varies among replicative sequences is evidence that it is of functional importance to the transposon. In the case of *Alu*, we also have potentially testable predictions about the changes in the effect of particular variations at particular points in time.

Plasmid mobilization assays support the contention that *Alu* replication operates under strong sequence constraint, although less extreme constraints than suggested by our analysis. Bennett et al. (11) identified 124 conserved positions in 282 sites among all elements shown to be replicative in a plasmid mobilization assay. We estimate that between 86 and 97 positions out of 175 non-CpG sites considered (excluding regions of difficult alignment) permit variation over the entire history of Alu, which suggests 126- 143 completely constrained positions in the 282 nucleotides at the core of *Alu.* Thus, around half of the constrained sites identified in our analysis may require a specific nucleotide for efficient replication by L1 proteins, similar to the estimate by Bennet (11). However, we find evidence that, due to changing sequence constraints, many sites that are compatible with replication at some point in time are restricted at other times, and on average only around 10% of sites are replication-compatible at any point in time. Plasmid assays take transposable element sequences out of their genomic contexts, and do not account for all possible sequence constraints; for example, in addition to affecting the efficiency with which *Alu* is retrotransposed by L1 proteins, the *Alu* sequence may also influence transcription levels (19) or host regulation. Thus, it is reasonable to presume that the plasmid mobilization assay identifies a minimum number of sites that would be constrained. It is also likely that selection is more sensitive than laboratory assays, particularly in the context of competition between expressed sequences.

Why might sequence constraints in *Alu* change over time? Both the primate host and the L1 transposon, which provides the machinery for *Alu* retrotransposition (20), likely experiences selective pressure to avoid parasitism by *Alu. Alu* elements impose substantial costs on the host by promoting genomic instability (21, 22) and by inserting in functional regions (23), and the host genome has numerous methods of regulating *Alu* activity (24–26). *Alu* retrotransposition activity may also impose costs on L1 through competition for proteins involved in retrotransposition (17). Thus, *Alu* may coevolve with the host, L1, or both in an evolutionary arms race, in which molecular changes reducing activity of the most successful classes of *Alu* drive compensatory changes in the *Alu* sequence. It is also possible that there are strong epistatic interactions within the *Alu* sequence that lead to changes in constraint over time.

## Methods

### Sequence Data

The sequence dataset was constructed by filtering the *Alu* annotations of the hg38 assembly of the human genome from RepeatMasker(28). Sequences outside of the 275- 325 bp length range were excluded to obtain a set of only full-length *Alu.*

### Ancestral Sequence Identification

We employed an iterative approach to identifying Alu ancestral sequences, taking additional sites into account in each round. In the first round, we considered the four sites with the highest frequency of non-consensus variants, excluding CpG sites. With one possible variant for each nucleotide type, this results in 4^4^=256 variant combinations.

Each round, we estimated two parameters for each variant combination. The parameter *F_i_* gives the *replicative frequency* of the variant combination. We define the replicative frequency of a combination as the expected number of surviving descendants of all ancestral sequences with that combination over the entire period of evolution of the TE family. The replicative frequency can be interpreted as a relative measure of the propensity of ancestral sequences with the combination to produce copies that survive to the present, integrated over the replicative lifetime of that combination. If *F_i_* = 0, that implies that there are no extant sequences descended from combination *i*.

The parameter *A_i_* is defined as the time point in which the ancestral sequences with combination *i* replicated. Here, we assume that the period of replicative activity of individual combinations was short relative to the evolution of the element, so that all descendants of combination *i* can be considered to be of age*A_i_*. Each round, we used a Gibbs sampler to estimate posterior distributions of *F_i_* and *A_i_* for all variant combinations, given our database of extant sequences (Supplementary Methods). Variant combinations that had at least 10% posterior probability of *F_i_* = 0 were then eliminated.

In each additional round, two more sites were considered, with four possible variants each for a total of 4^2^=16 variants. The 16 possible combinations of new variants were combined in all possible ways with all variant combinations remaining from the previous rounds (i.e., there were 16 times as many combinations considered in round *n* then there were remaining variant combinations after round *n*–1). This process was continued until all sites were considered.

## References

1. Chénais B, Caruso A, Hiard S, Casse N (2012) The impact of transposable elements on eukaryotic genomes: From genome size increase to genetic adaptation to stressful environments. Gene 509(1):7–15.

2. de Koning APJ, Gu W, Castoe TA, Batzer MA, Pollock DD (2011) Repetitive Elements May Comprise Over Two-Thirds of the Human Genome. PLoS Genet 7(12):094326.

3. Jurka J, Smith T (1988) A fundamental division in the Alu family of repeated sequences. Proc Natl Acad Sci 85(13):4775–4778.

4. Shen MR, Batzer MA, Deininger PL (1991) Evolution of the master Alu gene(s). J Mol Evol 33(4):311–320.

5. Deininger PL, Batzer MA, Hutchison III CA, Edgell MH (1992) Master genes in mammalian repetitive DNA amplification. Trends Genet 8(9):307–311.

6. Price AL, Eskin E, Pevzner PA (2004) Whole-genome analysis of Alu repeat elements reveals complex evolutionary history. Genome Res 14(11):2245–2252.

7. Wang J, et al. (2006) Whole genome computational comparative genomics: A fruitful approach for ascertaining Alu insertion polymorphisms. Gene 365:11–20.

8. Cordaux R, Hedges DJ, Batzer MA (2004) Retrotransposition of Alu elements: how many sources? Trends Genet 20(10):464–467.

9. Han K, et al. (2005) Under the genomic radar: The Stealth model of Alu amplification. Genome Res 15(5):655–664.

10. Wacholder AC, et al. (2014) Inference of Transposable Element Ancestry. PLoS Genet 10(8):094326.

11. Bennett EA, et al. (2008) Active Alu retrotransposons in the human genome. Genome Res 18(12)1875–1883.

12. Comeaux MS, Roy-Engel AM, Hedges DJ, Deininger PL (2009) Diverse cis factors controlling Alu retrotransposition: What causes Alu elements to die? Genome Res 19(4):545–555.

13. Price AL, Eskin E, Pevzner PA (2004) Whole-genome analysis of Alu repeat elements reveals complex evolutionary history. Genome Res 14(11):2245–2252.

14. Batzer MA, Deininger PL (2002) Alu repeats and human genomic diversity. Nat Rev Genet 3(5):370–379.

15. Price AL, Jones NC, Pevzner PA (2005) De novo identification of repeat families in large genomes. Bioinformatics 21(suppl 1):i351–i358.

16. Willard C, Nguyen HT, Schmid CW (1987) Existence of at least three distinct Alu subfamilies. J Mol Evol 26(3):180–186.

17. Slagel V, Flemington E, Traina-Dorge V, Bradshaw H, Deininger P (1987) Clustering and subfamily relationships of the Alu family in the human genome. Mol Biol Evol 4(1):19–29.

18. Cordaux R, Hedges DJ, Batzer MA (2004) Retrotransposition of Alu elements: how many sources? Trends Genet 20(10):464–467.

19. Liu W-M, Maraia RJ, Rubin CM, Schmid CW (1994) Alu transcripts: cytoplasmic localisation and regulation by DNA methylation. Nucleic Acids Res 22(6)1087–1095.

20. Dewannieux M, Esnault C, Heidmann T (2003) LINE-mediated retrotransposition of marked Alu sequences. Nat Genet 35(1):41–48.

21. Hedges DJ, Deininger PL (2007) Inviting Instability: Transposable elements, Doublestrand breaks, and the Maintenance of Genome Integrity. Mutat Res 616(1-2):46–59.

22. Ade C, Roy-Engel AM, Deininger PL (2013) Alu elements: an intrinsic source of human genome instability. Curr Opin Virol 3(6):639–645.

23. Deininger PL, Batzer MA (1999) Alu Repeats and Human Disease. Mol Genet Metab 67(3)183–193.

24. Liu WM, Schmid CW (1993) Proposed roles for DNA methylation in Alu transcriptional repression and mutational inactivation. Nucleic Acids Res 21(6):1351–1359.

25. Smalheiser NR, Torvik VI (2006) Alu elements within human mRNAs are probable microRNA targets. Trends Genet 22(10):532–536.

26. Obbard DJ, Gordon KHJ, Buck AH, Jiggins FM (2009) The evolution of RNAi as a defence against viruses and transposable elements. Philos Trans R Soc Lond B Biol Sci 364(1513):99–115.

27. Rouzic AL, Capy P (2006) Population Genetics Models of Competition Between Transposable Element Subfamilies. Genetics 174(2):785–793.

28. Smit A, Hubley R, Green P (2013) RepeatMasker 0pen-4.0.

